# Hippocampal astrocytes contribute to encoding context-specific aversive stimuli to regulate fear-related behavior

**DOI:** 10.64898/2026.05.22.727208

**Authors:** Daniela Sofia Abreu, Alexandra Veiga, Rafael M Jungmann, João Filipe Viana, Ana João Rodrigues, Carina Soares-Cunha, João Filipe Oliveira

**Affiliations:** Life and Health Sciences Research Institute (ICVS), School of Medicine, University of Minho, 4710-057 Braga, Portugal; ICVS/3B’s - PT Government Associate Laboratory, Braga/Guimarães, Portugal

## Abstract

In contextual fear conditioning, subjects learn to associate a neutral environment with an aversive stimulus and exhibit fear responses to that context, signaling danger. Neuronal activity in hippocampal-amygdala circuits has been shown extensively to underlie the conditioning and recall of contextual fear memories, yet the direct contribution of astrocytes to integrating stimulus and contextual information in this task remains unknown. In this study, we monitor astrocyte activity in real time in the hippocampus and simultaneously during performance in the contextual fear conditioning task. We find that shock-evoked astrocyte responses during conditioning predict the magnitude of freezing during recall in a context-specific manner. Importantly, astrocytic Ca^2+^ activity progressively precedes freezing onset specifically in the shock-associated context, indicating that hippocampal astrocytes contribute to encoding context-specific shock information. This work reveals an unprecedented contribution of astrocytes to the encoding of context-related information about aversive stimuli in fear-related behavior.

## Introduction

Animals develop fear responses to survive, especially in dangerous situations. Learned fear is a crucial survival mechanism because it allows animals to anticipate danger in neutral environments. The contextual fear conditioning (CFC) task is widely used to study fear learning in animal models, in which animals learn to associate a neutral setting with an aversive stimulus and exhibit fear-related freezing behaviors when danger is imminent (Maren *et al*., 2013). The processing of information encoding context- and fear-related stimuli requires the coordinated activity of neural circuits in the hippocampus and amygdala (Tovote *et al*., 2015). Although neuronal activity in these regions has been extensively characterized over the past decades, the advent of tools to monitor and manipulate activity in a cell-specific manner revealed unexpected roles for astrocytes in these circuits (Adamsky *et al*., 2018; Kol *et al*., 2020; Williamson *et al*., 2025; Dewa *et al*., 2025; Ghenissa *et al*., 2026).

Astrocytes are increasingly recognized as active participants in modulating brain circuits through dynamic intracellular calcium (Ca^2+^) signaling (Semyanov *et al*., 2020; Veiga *et al*., 2025). This dynamic Ca^2+^ activity modulates transmitter and neuromodulator release, ion homeostasis, and metabolic support, ultimately affecting neural circuit function (Hirrlinger & Nimmerjahn, 2022). Recent advances in genetically encoded Ca^2+^ indicators and *in vivo* imaging techniques have revolutionized our capacity to monitor the spatiotemporal complexities of astrocytic Ca^2+^ dynamics in the living brain, revealing their involvement in behavioral processing (Oliveira *et al*., 2015; Nagai *et al*., 2021).

Recent studies have confirmed that astrocytes in the hippocampus are involved in different phases of contextual fear conditioning. Specifically, astrocytes were shown to be involved in acquisition (Suthard *et al*., 2024), early (Adamsky *et al*., 2018; Li *et al*., 2020) and remote (Kol *et al*., 2020) recall, and fear extinction (Li *et al*., 2025). However, despite this progress, the acute contribution of hippocampal astrocytes to each phase, namely, to encoding and processing relevant information during the conditioning phase to modulate fear-related freezing behavior, is still unknown.

To address this critical knowledge gap, we implemented *in vivo* recording of dorsal hippocampus (dHIP) astrocytic Ca^2+^ dynamics throughout the contextual fear conditioning task. We developed an analysis pipeline that enables simultaneous detection and quantification of astrocyte Ca^2+^ signals during freezing behavior across acquisition and recall phases.

Our results suggest that dHIP astrocytes encode context-specific shock information during conditioning, thereby predicting and regulating fear-related freezing behavior.

## Results

### Astrocytic calcium activity is associated with the aversive stimulus in fear conditioning acquisition

We recorded Ca^2+^ activity in astrocytes of the dHIP of freely moving mice during CFC task using *in vivo* fiber photometry (**Fig. 1a**). Astrocytes expressed the Ca^2+^ indicator GfaABC1D-lck-jGCaMP8m in this region, as confirmed by immunohistochemistry. The expression was selective in astrocytes as confirmed by extensive co-expression with the GFAP marker and lack of co-expression with neuronal marker NeuN (**Fig. 1b**).

**Figure 1.**
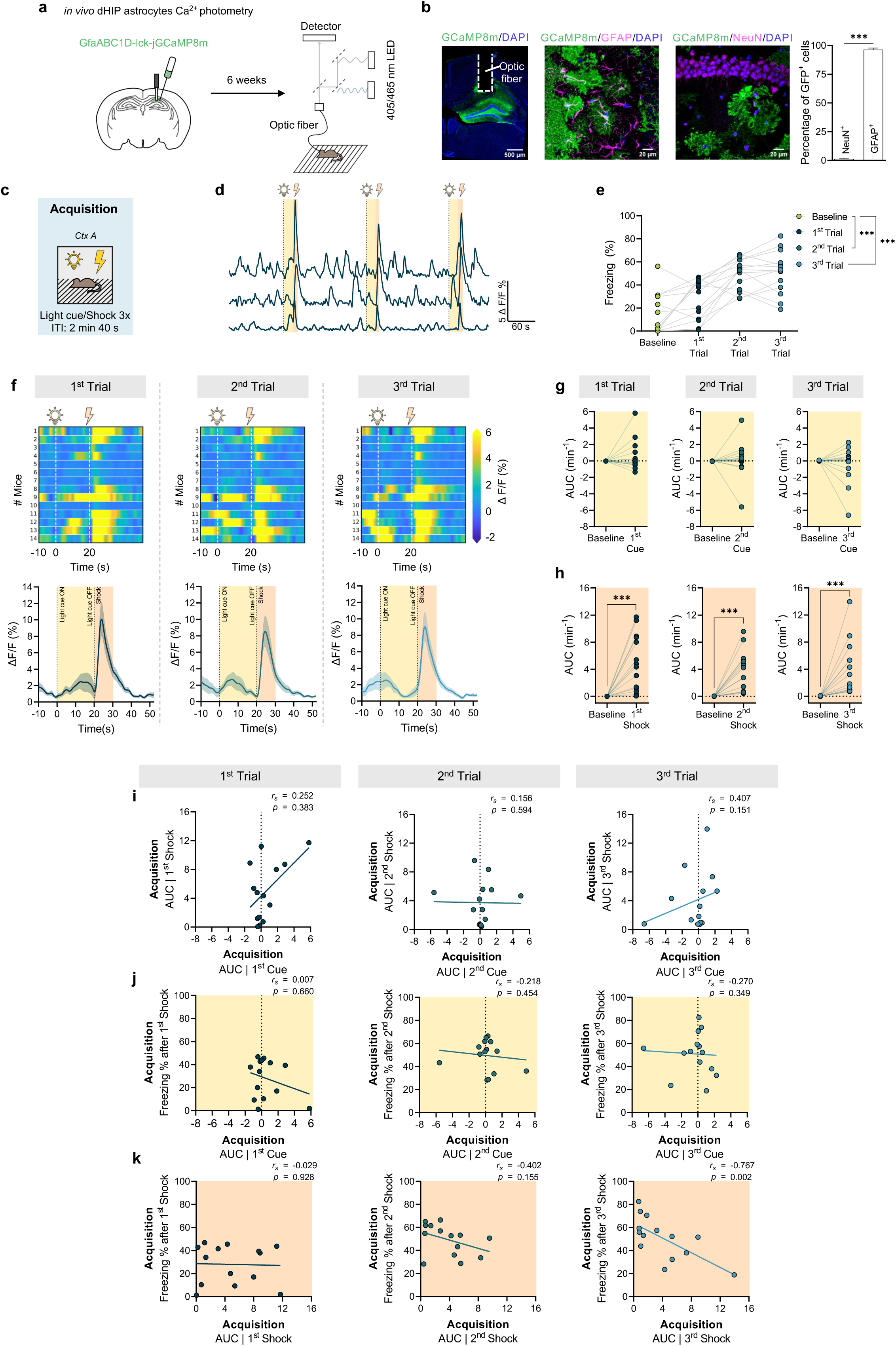
Shock-evoked calcium responses are associated with reduced immediate freezing after repeated shock exposure. **a**, In vivo Ca^2+^ fiber photometry in dorsal hippocampal (dHIP) astrocytes during contextual fear conditioning. **b**, Example fiber placement and GCaMP expression in astrocytes (GFAP) and neurons (NeuN). Scale bars, 500 μm (left) and 20 μm (middle and right). (*n* = 12 sections from 9 mice). **c**, Behavioral timeline (Cxt, context; ITI, Inter-Trial Interval). **d**, Representative **Δ**F/F traces from 3 mice during contextual fear acquisition. **e**, Freezing across testing trials (*n* = 14 mice). **f**, Population trace of CS and shock-related Ca^2+^ activity during acquisition, with three trial-wise heat maps above (*n* = 14 mice). **g**, AUC analysis of light cue (0–20 s) evoked Ca^2+^ activity during acquisition (*n* = 14 mice). **h**, AUC analysis of foot shock (20–30 s) evoked Ca^2+^ activity during acquisition (*n* = 14 mice). **i**, Correlation between the light cue-evoked Ca^2+^ and shock-evoked Ca^2+^, in each trial (*n* = 14 mice). **j**, Correlation between the light cue-evoked Ca^2+^ responses and freezing immediately after, across the three conditioning trials (*n* = 14 mice). **k**, Correlation between the foot shock-evoked Ca^2+^ responses and freezing immediately after, across the three conditioning trials (*n* = 14 mice). Data are presented as mean values ± SEM. *** p < 0.001.

Mice underwent a three-day CFC task comprising acquisition, recall with contextual discrimination, and cued recall. During the acquisition phase (**Fig. 1c**), in Context A (Ctx A) we delivered three pairs of a 20 s light cue with a 2 s foot shock and recorded dHIP astrocytic Ca^2+^ dynamics (**Fig. 1d**). Mice displayed the expected increase in periods of freezing behavior across the three conditioning trials (**Fig. 1e**), confirming effective acquisition of the conditioned response.

In each trial, we identified uneven Ca^2+^ responses in astrocytes during the period of light cue presentation (20 s) and strong Ca^2+^ responses after the foot shock, which lasted at least 10 s, despite the observed inter-animal variability (**Fig. 1f**). The integrated response (area under the curve, AUC) during the 20 s cue period did not differ from the baseline signal (**Fig. 1g**), suggesting that dHIP astrocytes did not significantly respond to the light cue in each trial. To assess temporal dynamics of astrocyte Ca^2+^ signals during cue presentation across trials, we analyzed the early (0-10 s) and late (10-20 s) epochs of the light cue response separately (**Fig. S1b**). We confirmed similar responses across trials in the early and late phases of cue presentations; thus, in subsequent analyses, we considered the full 20 s cue response. Finally, the signal AUC during the 20 s cue period was similar across the three light cue presentations (**Fig. S1c**).

Interestingly, the foot shock evoked a significant Ca^2+^ increase in dHIP astrocytes (**Fig. 1h**), in the 10 s after the shock in all trials. These foot shock-evoked responses were consistent across trials (**Fig. S1d**). The AUCs of cue-evoked responses were not correlated with foot shock response magnitude (**Fig. 1i**), suggesting that light cue and shock-evoked Ca^2+^ signals reflect independent mechanisms.

Next, we tested whether the magnitude of astrocytic responses to the light cue and shock correlated with behavioral fear expression on each trial. For each conditioning trial, we correlated cue- and shock-evoked Ca^2+^ signals with freezing behavior measured in the 1 min period following the stimulus. Cue-evoked Ca^2+^ responses did not correlate with freezing behavior measured in the 1 min period following the light cue on any trial (**Fig. 1j**). Surprisingly, shock-evoked Ca^2+^ responses along the 3 trials appear to align progressively with post-shock freezing, resulting in a significant negative correlation after the third shock, where mice with higher Ca^2+^ responses display lower freezing periods (**Fig. 1k**). Together, these data reveal an association with dHIP astrocytic Ca^2+^ responses to repeated exposure to an aversive stimulus (shock) specifically, but not to the light cue, with consequences for subsequent freezing behavior.

### Astrocytic activity during conditioning inversely correlates and predicts contextual fear expression during recall

To assess whether the progressive association between dHIP astrocyte Ca^2+^ responses and immediate freezing during acquisition would be maintained in the recall phase, we next correlated astrocytic Ca^2+^ activity during conditioning to subsequent contextual fear expression by the same animals.

24h after acquisition, mice were re-exposed to Ctx A (shock-paired context) for a 3 min contextual recall session, and 2 h later were placed in a novel neutral Context B (Ctx B; distinct from the conditioning environment, thus neither associated with the light cue, nor foot shock for a 3 min context discrimination session (**Fig. 2a**). As expected for this task, mice displayed increased freezing behavior in Ctx A indicating the establishment of contextual fear memory (**Fig. 2b**), and then a significant reduction of freezing to the new Ctx B, confirming a preserved context discrimination. We observed the typical inter-animal variability in freezing behavior in both contexts. Interestingly, we observed the establishment of a negative correlation between the extent of freezing behavior in Ctx A and astrocytic Ca^2+^ responses to the shock along the conditioning trials, which becomes very significant after trials 2 and 3 (**Fig. 2c**). Of note, this effect is highly context specific, as the astrocytic Ca^2+^ responses to shock are independent of freezing behavior on Ctx B (**Fig. 2d**). Moreover, this effect is specific to the Ca^2+^ responses to the shock, but not to the light cue and as there was no correlation between light cue responses during conditioning and freezing in either Ctx A (**Fig. S2a**) or Ctx B (**Fig. S2b**).

**Figure 2.**
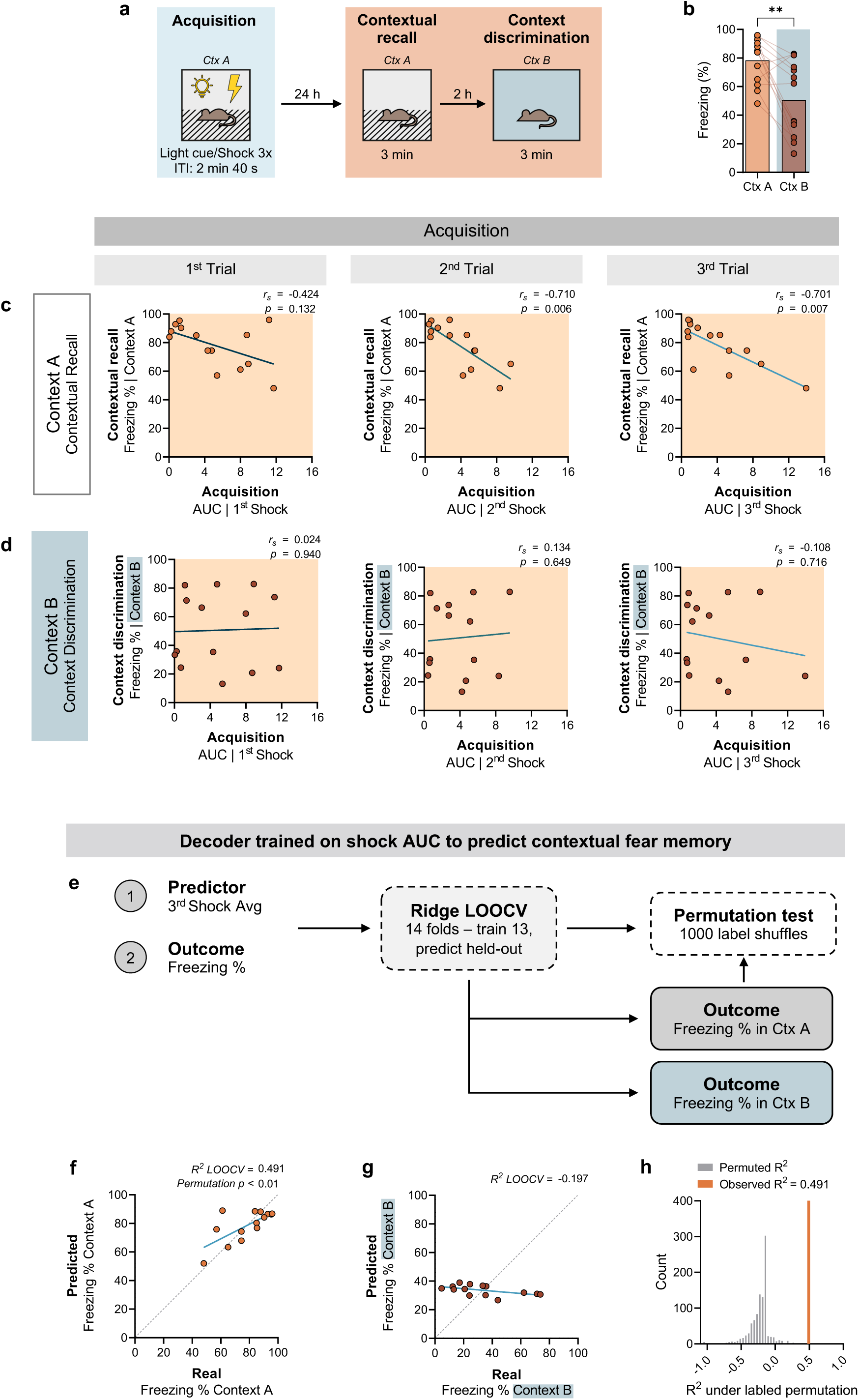
Astrocytic activity during conditioning inversely correlates and predicts contextual fear expression. **a**, Behavioral timeline (Cxt, context; ITI, Inter-Trial Interval). **b**, Freezing during contextual recall and context discrimination sessions (*n* = 14 mice). **c**, Correlation between the footshock-evoked Ca^2+^ responses across the three conditioning trials and freezing in Context A (*n* = 14 mice). **d**, Correlation between the footshock-evoked Ca^2+^ responses across the three conditioning trials and freezing in Context B (*n* = 14 mice). **e**, Schematic of Ridge regression predicting Context A and B tests freezing from the 3rd shock AUC. **f**, LOOCV predicted versus observed freezing in Context A (*n* = 14, each point represents one held-out animal). **g**, LOOCV predicted versus observed freezing in Context B (*n* = 14, each point represents one held-out animal). **h**, Observed R² (Context A, in orange) and the null distribution generated by repeating the full LOOCV pipeline on 1000 label-permuted datasets (n = 14 mice). Data are presented as mean values ± SEM. ** p < 0.01.

These data suggest that astrocytic Ca^2+^ responses to aversive conditioning stimuli may have predictive value for subsequent fear expression in a context-specific manner. To test this possibility, we applied Ridge regression with leave-one-mouse-out cross-validation (LOOCV) to the conditioning-shock AUC (**Fig. 2e**). The trial 3 AUC was used as the predictor feature. The Ridge LOOCV decoder explained 49.1% of Ctx A freezing variance (**Fig. 2f**). Decoder specificity was confirmed in Context B, where the same model failed to predict freezing above chance (**Fig. 2g**). The observed R^2^ significantly exceeded the shuffled null distribution generated by repeating the full LOOCV pipeline on 1000 label-permuted datasets (**Fig. 2h**), confirming robust and context-specific predictive power across mice.

These results suggest that dHIP astrocytes encode context-specific shock information during conditioning, which could predict and possibly regulate fear expression, as measured by freezing behavior, during recall.

### Astrocytic Ca^2+^ signals during the cue presentation do not encode cued freezing behavior

To confirm the specificity of the relationship between dHIP astrocyte Ca^2+^ and the shock versus the light cue to regulate freezing behavior, we next performed a cued test. 24 h after the contextual recall session, the same mice were presented with the 60 s light cue previously paired with foot shock during acquisition in Ctx B (neutral context, unpaired with shock or light cue) and (**Fig. 3a**). We analyzed Ca^2+^ dynamics during the light cue presentation and observed initial variable responses within the first 20 s and then a return to baseline (**Fig. 3b**). Therefore, we calculated AUC over this initial window, to dissect any potential relationship with freezing behavior and to avoid dilution by later baseline return. These initial Ca^2+^ signals did not differ from their respective baselines (**Fig. 3c**), consistent with the observations during conditioning (**Fig. 1g**). Nevertheless, we quantified freezing responses in the initial 20 s and the full 60 s period to assess any correlation with inter-animal variability. The initial astrocytic Ca^2+^ responses during light cue presentation failed to correlate with the inter-animal differences in freezing behavior measured during the first 20 s (**Fig. 3d**) or across the full 60 s period (**Fig. 3e**).

**Figure 3.**
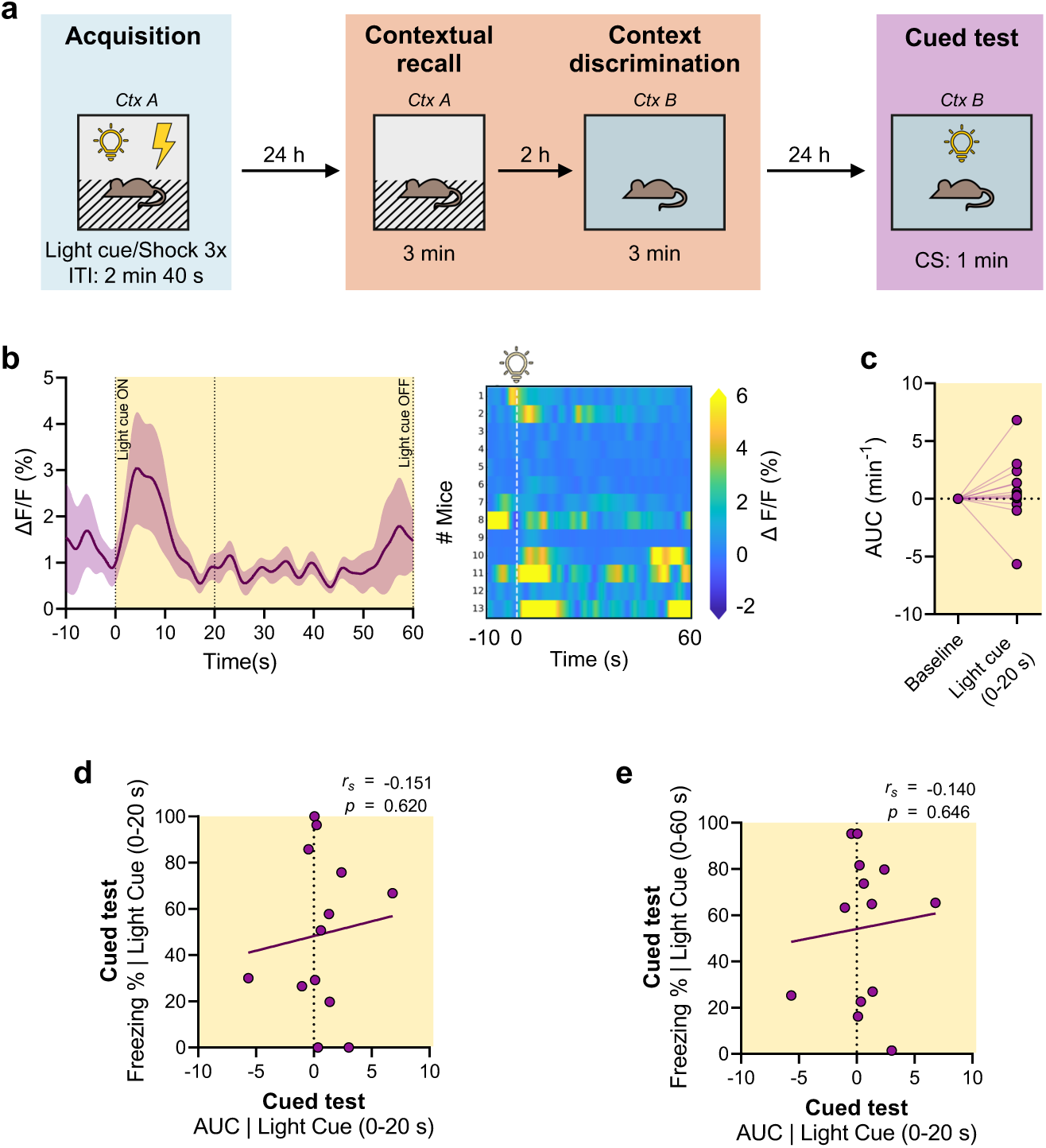
Astrocytic calcium responses in the dHIP do not encode cued fear expression. **a**, Behavioral timeline (Cxt, context; ITI, Inter-Trial Interval). **b**, Population trace of cued CS-related Ca^2+^ activity during cued test with heat map of the right side (*n* = 13 mice). **c**, AUC analysis of light cue (0–20 s) Ca^2+^ activity during cued test (*n* = 13 mice). **d**, Correlation between the light cue-evoked Ca^2+^ responses during the first 20 s of the cued test, and the freezing during the first 20 s of light cue exposure during the cued test (*n* = 13 mice). **e**, Correlation between the light cue-evoked Ca^2+^ responses during the first 20 s of the cued test, and the freezing during the entire 60 s of light cue exposure during the cued test test (*n* = 13 mice). Data are presented as mean values ± SEM.

Together, these data show that neither concurrent nor prior astrocytic Ca^2+^ responses to the light cue predict cued fear expression, suggesting that dHIP astrocytes specifically encode shock information rather than cue information.

Until now, our data suggest that astrocytes in the dHIP encode context-specific information about shock, which is required to regulate the extent of mouse freezing.

To confirm this hypothesis, we conducted a series of experiments to assess whether astrocyte Ca^2+^ activity is required to regulate the extent of fear expression and whether astrocytes encode information at the initiation or termination of freezing behavior.

### Astrocytic Ca^2+^ signaling is required to regulate the extent of fear-related freezing behavior

To test whether dHIP astrocytic Ca^2+^ activity is required for regulating fear expression, we next assessed *IP_3_R2* knockout (*IP_3_R2 KO*) mice in the same CFC task. These mice were shown by different laboratories to lack somatic Ca^2+^ signaling specifically in astrocytes (Petravicz *et al*., 2008; Navarrete *et al*., 2012; Srinivasan *et al*., 2015), but to display normal acquisition of developmental milestones and adult motor coordination, strength, and neurological reflexes (Guerra-Gomes *et al*., 2021), confirming their suitability for adult behavior assessment. We used the same viral approach to drive the expression of the genetically encoded Ca^2+^ indicator (jGCaMP8m) under the astrocytic promoter (GfaABC1D) in astrocytes of the dHIP and performed *in vivo* fiber photometry (**Fig. 4a**). We observed a complete abolishment of Ca^2+^ activity in dHIP astrocytes (**Fig. 4b**), and a marked reduction of spontaneous Ca^2+^ activity (**Fig. 4c**).

**Figure 4.**
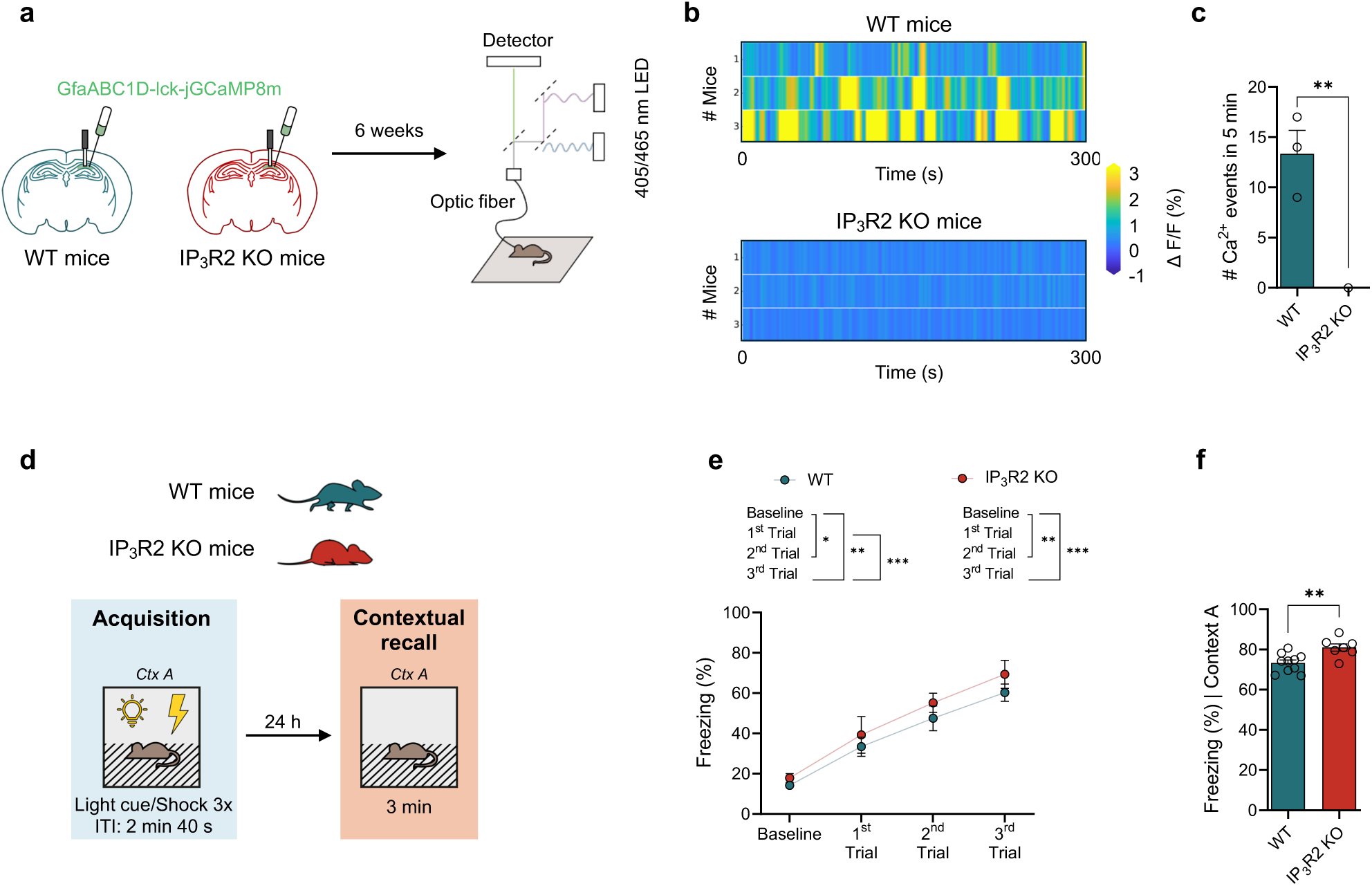
Astrocytic calcium signaling is required to regulate the extent of fear-related freezing behavior. **a**, In vivo Ca^2+^ fiber photometry in dorsal hippocampal astrocytes of WT and IP_3_R2 KO mice. **b**, Heatmap of population Ca^2+^ activity during open field test (*n* = 3 WT mice, *n* = 3 IP_3_R2 KO mice). **c**, Quantification of Ca^2+^ event frequency per mouse during the entire open field session (*n* = 3 WT mice, *n* = 3 IP_3_R2 KO mice). **d**, Behavioral timeline (Cxt, context; ITI, Inter-Trial Interval). **e**, Freezing across the three acquisition trials of WT and IP_3_R2 KO mice (*n* = 13 WT mice, *n* = 7 IP_3_R2 KO mice). **f**, Freezing during contextual recall of WT and IP_3_R2 KO mice (*n* = 13 WT mice, *n* = 7 IP_3_R2 KO mice). Data are presented as mean values ± SEM. ** p < 0.01, *** p < 0.001.

We then examined the consequences of this lack of astrocytic Ca^2+^ activity in the same CFC task (**Fig. 4d**). During acquisition, as expected, mice increase the freezing responses to each shock from trial to trial, in a similar manner between groups, indicating similar fear conditioning (**Fig. 4e**). Interestingly, during the recall in Ctx A, *IP_3_R2 KO* mice exhibited higher freezing behavior than *WT* littermate controls (**Fig. 4f**), suggesting that the astrocytic Ca^2+^ is required for the regulation of the extent of freezing behavior, suggesting an involvement in fear memory processing.

Overall, these results suggest that astrocyte Ca^2+^ activity is an integral component of fear memory expression, as it is required to regulate the extent of freezing behavior in the CFC task.

### Astrocytic Ca^2+^ activity preceding freezing onset regulates the extent of fear expression

Our data suggests that dHIP astrocytes encode context-specific information about shock, which is required to regulate the extent of mouse freezing. Thus, we conducted a detailed analysis of the temporal dynamics of astrocytic Ca^2+^ signals to assess whether astrocytes encode information at the initiation or termination of freezing behavior. To enable this analysis, we developed a Python routine to align astrocyte Ca^2+^ signals to freezing onset (**Fig. 5a**) and offset (**Fig. 5b**) events. We analyzed freezing events within the 1 min window following each foot shock delivery during conditioning in the acquisition phase. We observed the progressive appearance of an astrocyte Ca^2+^ signal at the freezing onset across all three trials (**Fig. 5c-e**). Moreover, we observed a decrease in astrocytic Ca^2+^ terminating after the freezing offset (**Fig. 5f-h**). For each animal and trial, we quantified the slope of the Ca^2+^ trace in two epochs: 2 s before (-2 to 0 s) and 2 s after (0 to +2 s) the transition. This quantification allowed us to confirm that the pre-onset slope became progressively steeper across trials and was significantly higher on the third shock (**Fig. 5i**). This effect is specific to the pre-onset period, as the negative post-onset slope remained stable across trials (**Fig. 5j**). In contrast, astrocyte Ca^2+^ signal slopes around both pre-offset (**Fig. 5k**) and post-offset (**Fig. 5l**) of freezing behavior remain stable across the conditioning trials.

**Figure 5.**
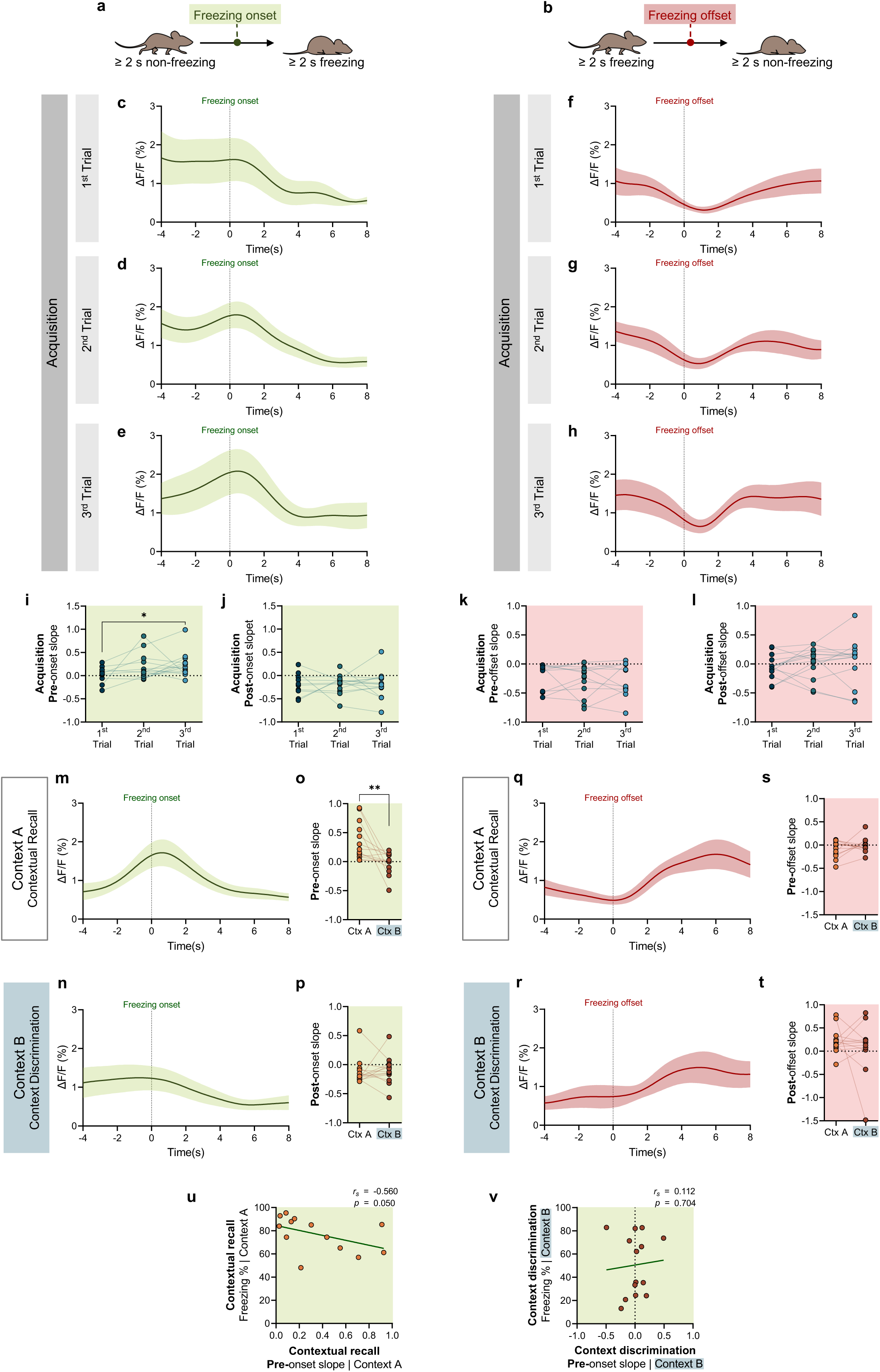
Astrocytic calcium activity preceding freezing onset regulates the extent of fear expression. **a**, Freezing onset (initiation) scheme. **b**, Freezing offset (termination) scheme. **c**, Astrocytic Ca^2+^ traces during freezing onset, during the 1^st^ Trial of acquisition (*n* = 10 mice with valid freezing events). **d**, Astrocytic Ca^2+^ traces during freezing onset, during the 2^nd^ Trial of acquisition (*n* = 14 mice with valid freezing events). **e**, Astrocytic Ca^2+^ traces during freezing onset, during the 3^rd^ Trial of acquisition (*n* = 14 mice with valid freezing events). **f**, Astrocytic Ca^2+^ traces during freezing offset, during the 1^st^ Trial of acquisition (*n* = 10 mice with valid freezing events). **g**, Astrocytic Ca^2+^ traces during freezing offset, during the 2^nd^ Trial of acquisition (*n* = 14 mice with valid freezing events). **h**, Astrocytic Ca^2+^ traces during freezing offset, during the 3^rd^ Trial of acquisition (*n* = 12 mice with valid freezing events). **i**, Slope analysis -2 to 0 s before freezing onset during acquisition, across the three trials (1^st^ Trial *n* = 10 mice with valid freezing; 2^nd^ Trial *n* = 14 mice with valid freezing events; 3^rd^ Trial *n* = 14 mice with valid freezing events). **j**, Slope analysis 0 to +2 s after freezing onset during acquisition, across the three trials (1^st^ Trial *n* = 10 mice with valid freezing; 2^nd^ Trial *n* = 14 mice with valid freezing events; 3^rd^ Trial *n* = 14 mice with valid freezing events). **k**, Slope analysis -2 to 0 s before freezing offset during acquisition, across the three trials (1^st^ Trial *n* = 10 mice with valid freezing; 2^nd^ Trial *n* = 14 mice with valid freezing events; 3^rd^ Trial *n* = 12 mice with valid freezing events). **l**, Slope analysis 0 to +2 s after freezing offset during acquisition, across the three trials (1^st^ Trial *n* = 10 mice with valid freezing; 2^nd^ Trial *n* = 14 mice with valid freezing events; 3^rd^ Trial *n* = 12 mice with valid freezing events). **m**, Astrocytic Ca^2+^ traces during freezing events onset, during contextual recall session, in Context A (*n* = 13 mice with valid freezing events). **n**, Astrocytic Ca^2+^ traces during freezing events onset, during context discrimination session, in Context B (*n* = 13 mice with valid freezing events). **o**, Comparison of slope -2 to 0 s before freezing onset in Context A and Context B (*n* = 13 mice). **p**, Comparison of slope -0 to +2 s after freezing onset in Context A and Context B (*n* = 13 mice). **q**, Astrocytic Ca^2+^ traces during freezing events offset, during contextual recall session, in Context A (*n* = 13 mice with valid freezing events). **r**, Astrocytic Ca^2+^ traces during freezing events offset, during context discrimination session, in Context B (*n* = 13 mice with valid freezing events). **s**, Comparison of slope -2 to 0 s before freezing offset in Context A and Context B (*n* = 13 mice). **t**, Comparison of slope -0 to +2 s after freezing offset in Context A and Context B (*n* = 13 mice). **u**, Correlation between slope - 2 to 0 s before freezing onset in Context A and freezing in Context A (*n* = 13 mice). **v**, Correlation between slope -2 to 0 s before freezing onset in Context B and freezing in Context B (*n* = 13 mice). Data are presented as mean values ± SEM. * p < 0.05, ** p < 0.01.

Together, these results suggest that dHIP astrocytic Ca^2+^ activity selectively increases pre-freezing onset and scales with repeated shock exposure, while Ca^2+^ dynamics around freezing offset remain stable across trials.

Given the progressive pre-onset Ca^2+^ ramp observed during conditioning, we next investigated whether this activity pattern is preserved in a context-specific manner during fear memory expression. To this end, we aligned dHIP astrocytic Ca^2+^ activity to freezing onset and offset during contextual recall in Ctx A and during the context discrimination session in Ctx B. In Ctx A, we observed a clear astrocytic Ca^2+^ signal preceding the freezing onset characterized by a positive pre-onset slope (**Fig. 5m**), recapitulating the pattern observed during conditioning. Interestingly, in Ctx B, this signal was absent, resulting in a pre-onset slope close to zero (**Fig. 5n**). Consistently, the pre-onset Ca^2+^ signals were significantly larger in Ctx A than in Ctx B (**Fig. 5o**), whereas post-onset slopes did not differ between contexts (**Fig. 5p**). By contrast, astrocytic Ca2+ signals were similar at freezing offset in Ctx A (**Fig. 5q**) and Ctx B (**Fig. 5r**), as given by equivalent Ca^2+^ slopes pre- and post-freezing termination (**Fig. 5s, t**).

Together, these findings suggest that astrocytes selectively encode information to initiate context-specific freezing, possibly modulating the extent of fear memory expression. This idea is corroborated by the fact that mice that have steeper pre-freezing astrocytic Ca^2+^ signals display shorter freezing behavior in Ctx A (**Fig. 5u**). Moreover, this effect is context-specific, as pre-onset Ca^2+^ signal slope fails to correlate with freezing extent in Ctx B (**Fig. 5v**). These findings support the idea that astrocytic Ca^2+^ activity preceding freezing onset predicts the extent of fear memory expression in a context-dependent manner.

## Discussion

In this study, we selectively monitored Ca^2+^ dynamics in dHIP astrocytes during contextual fear conditioning and found that shock-evoked responses inversely predicted the magnitude of later freezing specifically in the shock-paired context. Importantly, we also found that astrocytic Ca^2+^ elevations specifically preceded freezing onset in the shock-associated context, and that steeper pre-freezing ramps are associated with reduced contextual fear memory expression. Overall, the progressive emergence of these effects suggests that dHIP astrocytes encode context-specific shock information during conditioning, thereby inversely predicting and regulating the initiation and the subsequent extent of fear memory expression.

Next, we will discuss our main findings supporting the conclusions in light of the existing literature, namely the shock- versus cue specificity of hippocampal astrocyte Ca^2+^ signals, the progressiveness of the relationship of shock-evoked astrocyte Ca^2+^ and the extent of freezing behavior, the meaning of this context-specific negative correlation, and finally how this type of information encoding may constitute a complementary mechanism of fear-related memory.

In this study, we analyzed astrocytic Ca^2+^ responses to the unconditioned foot shock and to the conditioned light cue. We observed that only shock-evoked Ca^2+^ signals in dHIP astrocytes correlated with behavioral output, whereas responses during light cue presentation remained independent of freezing behavior across trials in days 1 and 3 of the CFC task. These observations are consistent with previous evidence indicating that the hippocampus is involved in contextual encoding of fear memory, whereas discrete conditioning cues are represented in the amygdala, suggesting that astrocytes in the dHIP follow that rule, participating in the contextual and state-dependent modulation rather than directly integrating each light cue (Maren *et al*., 2013; Plas *et al*., 2024).

Interestingly, the specific link between shock-evoked dHIP astrocytic Ca^2+^ and reduced contextual fear expression is inversely associated. This association is supported by our findings in *IP_3_R2 KO* mice, which lack astrocytic somatic Ca^2+^ activity and display increased fear memory. This observation is also consistent with a previous study showing that astrocytic Ca^2+^ reduction in dorsal CA1 via hPMCA2w/b (Ca^2+^-extruding pumps) expression increased contextual freezing without affecting cued fear (Li *et al*., 2020). This study also showed the opposite effect: optogenetic activation of CA1 astrocytes in transgenic GFAP-ChR2 rats reduced contextual, but not cued, fear memory.

The progressive negative correlation between shock-evoked astrocytic Ca^2+^ activity and post-shock freezing across conditioning trials suggests that dHIP astrocytes are engaged to encode the aversive experience, rather than reflecting an immediate linear consequence. The next question was whether astrocytic signaling temporally antecipates fear processing. Here, our synchronized *in vivo* recording of astrocytic Ca^2+^ activity during freezing episodes provides this missing piece. Rather than co-varying with freezing behavior, dHIP astrocytic Ca^2+^ increased approximately 1 s before freezing onset. Of note, this anticipatory Ca^2+^ increase emerged progressively during conditioning with repeated aversive exposure, becoming consistently detectable only after the third foot shock, suggesting that it reflects a learned association (suggesting a form of information encoding) rather than an innate response to threat. We believe that while neurons in these circuits are performing their information encoding on the millisecond timescale, astrocytes might read neuronal communication, interpret the context, and update the circuit with network state information (Murphy-Royal *et al*., 2023) across conditioning trials, enabling progressive neuronal encoding in the next trial.

Again, this progressive encoding of astrocyte Ca^2+^ activity and freezing onset is context-specific, since the anticipatory Ca^2+^ increase preceded freezing in the aversive context but was absent during freezing episodes in a neutral, non-aversive environment, indicating that, also here, dHIP astrocytes are not encoding the motor act of freezing itself but rather its contextually appropriate expression. The observed link between hippocampal astrocytic Ca^2+^ and freezing behavior appears to have different kinetics from that relating astrocytic responses upon movement initiation in the somatosensory cortex (Fedotova *et al*., 2023). Nevertheless, we observe an increase in Ca^2+^ activity on the second time scale after freezing offset in both contexts, which could be associated with movement initiation.

The ramping increase preceding freezing onset, followed by a signal decay to baseline, recapitulates Ca^2+^ dynamics reported in basolateral amygdala astrocytes during contextual fear acquisition and expression, where astrocytic activity becomes time-locked to freezing initiation and termination (Suthard *et al*., 2023, 2024). Together, these data indicate that hippocampal astrocytes do not passively track generic immobility but actively encode the transition into contextually appropriate fear states through Ca^2+^ signaling that anticipates behavioral output.

The predictive relationship between dHIP astrocytic activity during acquisition and contextual fear expression at recall suggests that modulation of astrocyte Ca^2+^ signals integrates the neuronal mechanisms involved in fear memory processing in a context-dependent manner. Together, these properties, temporal anticipation of behavior, emergence with learning, alignment during memory recall with context specificity, and influence in freezing initiation, suggest that dHIP astrocytic Ca^2+^ participates in the encoding and subsequent readout of the contextual fear memory. This is consistent with the consolidation-phase role previously described (Li *et al*., 2020)and with the broader framework of learning-associated astrocyte recruitment recently described in the hippocampus (Williamson *et al*., 2025), accumbens (Serra *et al*., 2025), or amygdala (Dewa *et al*., 2025) and elegantly discussed (Sánchez Romero & Navarrete, 2026)). Whether this anticipatory signal actively reflects the recruitment of the astrocyte by the contextual memory engram prior to behavioral manifestation remains an open question. Also open is the relationship to the observation that modulation of astrocyte activity using DREADDs causes alternative responses (Adamsky *et al*., 2018; Kol *et al*., 2020). Indeed, these studies confirm that hippocampal astrocytes are involved in fear conditioning, as their manipulation triggers robust behavioral responses. Nevertheless, the mechanism underlying these effects needs to be clarified, as a linear Gq-DREADD increase in activation-driven fine astrocyte Ca^2+^ is unlikely based on recent data. The conventional assumption underlying chemogenetic studies (originally developed to manipulate neurons) has been that activation of hM3Dq (Gq-DREADD) elevates astrocytic Ca^2+^ levels, whereas activation of hM4Di (Gi-DREADD) suppresses it, providing bidirectional causal control over astrocytic Ca^2+^ signaling. However, recent work (Bukalo *et al*., 2026) documented the temporal Ca^2+^ profile following hM3Dq and hM4Di manipulations, and observed that neither manipulation produces the simple, sustained scalar shift in Ca^2+^ that multiple frameworks assumed *in vivo*. Following hM3Dq activation, astrocytic Ca^2+^ exhibits a brief surge lasting approximately 10 min, after which activity falls below baseline and remains suppressed for at least 2 h. Critically, this prolonged suppression reduces the foot shock-evoked astrocytic Ca^2+^ responses. Conversely, rather than suppressing Ca^2+^, hM4Di activation increases the cumulative frequency of spontaneous Ca^2+^ transients and enhances stimulus-evoked responses. These observations are in line with other recent studies using these tools, which describe maximal astrocyte Ca^2+^ during prolonged periods of CNO bioavailability, which render cells dysfunctional (Delepine *et al*., 2023; Miguel-Quesada *et al*., 2023).

The innovative live imaging and synchronization of astrocyte activity in the dHIP throughout the performance of the contextual fear conditioning task enabled the dissection of a surprising temporal relationship between astrocyte activity and aversive stimulus conditioning and consequent fear responses. Our data support the view that hippocampal astrocytes contribute to the encoding of context-specific shock information during conditioning, thereby predicting and regulating the initiation and subsequent extent of fear memory expression.

## Methods

### Subjects

All procedures involving mice were conducted in accordance with the European Directive 2010/63/EU on the protection of animals used for scientific purposes. Experimental protocols were approved by the Ethics Committee of the Life and Health Sciences Research Institute (ICVS) and by the national authority for animal experimentation, Direção-Geral de Alimentação e Veterinária (DGAV; approval reference DGAV 023838/2019 and DGAV 34850/24-S). Health monitoring followed FELASA recommendations, and all personnel and facilities were certified by DGAV.

Mice were housed in groups of 2-6 per cage under controlled conditions (22 ± 1 °C, 55% relative humidity, 12 h light/dark cycle) with *ad libitum* access to food and water. In this study, we employed male C57BL/6J mice, as well as wild-type (*WT*) and respective homozygous *IP_3_R2* knockout (KO) littermates, aged 8-12 weeks.

*WT* and *IP_3_R2 KO* mice were maintained on a C57BL/6J background, as previously described (Guerra-Gomes *et al*., 2021). Founder breeding pairs were provided by Prof. Alfonso Araque (University of Minnesota, USA) (Navarrete *et al*., 2012), under a material transfer agreement with Prof. Ju Chen (University of California, San Diego, USA) (Li *et al*., 2005).

Mice were ear-tagged at weaning (postnatal day 21), and the tags remained unaltered throughout the study to enable animal identification and blind data analysis. Genotyping was performed by PCR from ear biopsies using *WT*-specific primers (forward: ACCCTGATGAGGGAAGGTCT; reverse: ATCGATTCATAGGGCACACC) and mutant allele-specific primers (neo-specific forward: AATGGGCTGACCGCTTCCTCGT; reverse: TCTGAGAGTGCCTGGCTTTT), as previously described (Li *et al*., 2005).

### Surgeries

All surgical procedures were performed under aseptic conditions. Mice were anesthetized with sevoflurane (2-3% in 1.0-1.5 L min^-1^ oxygen) delivered via a calibrated vaporizer and positioned on a stereotaxic frame (Model 940, David Kopf Instruments, USA). Body temperature was maintained at 36 °C using a heating pad (#53850-MM, Stoelting). After surgery, animals were allowed to recover in their home cages under a heating lamp until full ambulation.

Pre- and post-operative analgesia were administered with buprenorphine (0.05 mg kg^-1^, intraperitoneal; Bupaq, Richter Pharma, Austria). Postoperative monitoring occurred daily for up to 3 days, with additional buprenorphine administration if signs of discomfort were observed.

Following scalp shaving and disinfection, an incision was made to expose the skull. Bregma and lambda were aligned in the same horizontal plane, along the left-right axis. A small craniotomy was drilled using a high-speed drill (#5144, Stoelting) at stereotaxic coordinates targeting the dHIP, from bregma (Franklin & Paxinos, 2007): 1.8 mm anteroposterior, ± 1.3 mm mediolateral, and -1.3 mm dorsoventral.

For photometry experiments, mice received bilateral injections of 1 µL of AAV1 GfaABC1D lck jGCaMP8m (Addgene #176760; titer = 5.5 × 10^12^ genome copies mL^-1^). Viral delivery was performed using a syringe pump (Quintessential Stereotaxic Injector, Stoelting). Connected to a 10 µL Hamilton syringe (#80330) and a 30 G needle (#7803-07). The injection rate was maintained at 100 nL min^-1^. After infusion, the needle was kept in place for 5 min to allow viral diffusion before slow withdrawal. Optical fibers were implanted at the same coordinates, adjusted to a dorsoventral position of -1.25 mm. Implants consisted of a Ø1.25 mm diameter zirconia ferrule housing a Ø400 µm core multimode optical fiber with a numerical aperture of 0.50 (ThorLabs, USA). Ferrules were secured to the skull using dental cement (C&B Kit, Sun Medical, Japan).

### Contextual Fear Conditioning

Behavioral experiments were performed during the light phase. Mice were handled for 10 min daily for at least 10 days before testing. 5 days before behavioral experiments, animals were habituated to a patch cable for 5 min per day in an empty home cage. Behavioral sessions were conducted by a non-blinded investigator, whereas data analysis was performed blind to genotype.

The contextual fear conditioning (CFC) protocol consisted of three consecutive daily sessions. On Day 1 (acquisition), mice were tested in Context A (Ctx A; 29.5 × 25.0 × 11.5 cm; Med Associates Inc., San Diego, CA, USA), comprising a stainless-steel shock grid floor. Each mouse was placed in Ctx A and allowed to freely explore for 160 s before receiving three light cue-shock pairings. Each pairing consisted of a 20 s light cue immediately followed by a 2 s foot shock (0.5 mA). Inter-trial intervals were 160 s. The first CS presentation was preceded by a 1 s TTL pulse to synchronize photometry acquisition.

On Day 2 (contextual recall), 24 h after acquisition, mice were re-exposed to Ctx A for 3 min without any stimulus. 2 h later, they were placed in a novel Context B (Ctx B; 21.5 × 19.5 × 17.0 cm; Med Associates Inc.) for 3 min to assess context discrimination. Ctx B differed from Ctx A by visual and olfactory cues (walls lined with blue cardboard, vanilla odor). The experimenter changed lab coats and gloves between contexts to minimize the transfer of cues. In both contextual recall and context discrimination sessions, a 1 s TTL pulse preceded the activation of a red LED, visible in the top-view video feed, to enable temporal synchronization between CFC behavior and recorded photometry data.

On Day 3 (cued recall), mice were re-exposed to Ctx B, and the conditioned light cue was presented for 60 s. A 1 s TTL pulse preceded the light cue onset for synchronization with photometry recordings. Mice were returned to their home cages 30 s after cue termination. Between trials, Ctx A was cleaned with 10% ethanol, and Ctx B was cleaned with distilled water scented with vanilla.

Behavioral sessions were conducted in sound-attenuating acrylic enclosures (63.5 × 43.2 × 44.5 cm; Med Associates Inc.) equipped with a dim back-mounted light source. Behavior was recorded using the OBS software via a ceiling-mounted digital video camera. Photometry recordings were synchronized with behavioral recordings via TTL pulses (details to follow in the “Fiber photometry recordings” section).

### Fiber photometry recordings

Astrocytic Ca^2+^ signals were recorded using a dual-excitation fiber photometry system (Doric Lenses, Quebec, Canada) controlled by Doric Studio software. A patch cable (Ø400 µm core and 0.5 NA, with 1m, Doric Lenses, Quebec, Canada) was coupled to the implanted optical fiber via an ADAL3 interconnect for Ø 1.25 mm ferrules (ThorLabs, Newton, NJ, USA).

Excitation light was produced by two LEDs (models CLED_465 and CLED_405; Doric Lenses) driven by a dual-channel LED driver (Doric Lenses). Emission and excitation light were combined and spectrally separated through a minicube (Doric Lenses). The 465 nm excitation wavelength selectively excited GCaMP8m fluorescence, whereas the 405 nm channel served as the isosbestic control to correct for movement and photobleaching artifacts.

The emitted fluorescence signal was transmitted through the same optical fiber, separated from excitation light by integrated dichroic filters, and converted to electrical signals by a high-sensitivity photoreceiver (model 2151, Newport Corporation) equipped with a lensed FC adapter (Doric Lenses). A TTL synchronization input on a separate channel was used to align photometry recordings with behavioral events and video tracking. Fiber photometry data were exported as CSV files. Each file contained time-stamped voltage signals from a Ca^2+^-sensitive channel (GCaMP8m, ∼465 nm excitation), an isosbestic control channel (tdTomato, ∼405 nm excitation), a digital TTL channel, and a time column.

All analyses were performed in Python 3.10.18 (Anaconda distribution, 64-bit, Windows) using the following libraries: NumPy 1.26.4, SciPy 1.15.3, pandas 2.3.1, Matplotlib 3.10.0, seaborn 0.13.2, and scikit-learn 1.7.2. Data serialization used Python’s pickle module. Analysis scripts were developed with the assistance of Claude (Anthropic, claude.sonnet-4-6), an AI-based coding assistant. All code was verified by the authors against the described methodology before use. Data input/output relied on the pandas CSV reader and the pickle module for serialized array storage. All operations were applied sequentially per recording file in a deterministic, single-pass pipeline; no stochastic operations or random seeds were required.

### TTL pulse detection

TTL offset times were identified on the original, high-resolution signal before downsampling to prevent edge blurring introduced by resampling. Upward transitions (0→1) in the binary digital channel were detected by locating indices where the discrete first difference equaled 1. The time stamp of the sample immediately following each transition was recorded as the TTL offset time.

### Downsampling

To ensure a consistent temporal resolution across recordings, all signals (GCaMP, isosbestic control, and TTL) were downsampled to 100 Hz (10 ms intervals) using the pandas DataFrame resampling function. Timestamps were computed relative to the first sample of each recording. The resulting sampling rate was verified numerically as the reciprocal of the mean inter-sample interval.

### Temporal cropping and alignment to behavioral events

A median filter with a 5-sample kernel was applied to both GCaMP and isosbestic signals to remove high-frequency electrical noise.

All recordings were temporally cropped to begin 120 s before the first TTL pulse (light cue onset). Following cropping, the time axis was re-zeroed such that t = 0 s corresponds exactly to the first light cue onset, creating a standardized reference frame where negative times represent pre-first light cue baseline activity and positive times represent fear conditioning event-locked responses. This approach provides a 2 min window for baseline characterization while maintaining precise alignment of astrocytic activity to behavioral events across all subjects.

### ΔF/F calculation

Fiber photometry data were initially processed using a free open-source package ‘Guided Photometry Analysis in Python’ (GuPPy (Sherathiya *et al*., 2021)) and as previously described in (Deseyve *et al*., 2024). The isosbestic signal was fit to the GCaMP8m signal using ordinary least squares linear regression. Fluorescence changes were quantified using the standard ΔF/F metric, defined as:

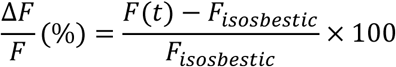

where *F*(*t*) is the Ca^2+^-dependent fluorescence of GCaMP, at time *t*, and *F_isosbestic_*represents the isosbestic channel fluorescence.

### Correction of motion artifact, photobleaching, and baseline drift

Motion artifacts were detected and corrected in the ΔF/F signal using a derivative-based median absolute deviation (MAD) approach (Serra *et al*., 2025). MAD estimates the typical variability of the derivative signal while remaining minimally influenced by extreme values. Time points at which the rate of change exceeded four times the MAD relative to the median were classified as motion artifacts. The identified artifact-contaminated samples (above 4×MAD) were then corrected using linear interpolation between the nearest unaffected data points flanking the artifact. This approach maintains signal continuity and preserves the overall trajectory of the recording while eliminating artifact-induced distortions.

To correct slow baseline drift due to photobleaching and mechanical changes in fiber coupling, a sliding-window percentile-based baseline estimation was applied to the ΔF/F signal after motion artifact removal (Serra *et al*., 2025). A rolling window of width equal to 5% of the total signal length in samples was used (minimum window size: 50 samples).

For each window position, the 10th percentile of signal amplitudes within that window was computed, producing a continuous trace that tracks the slowly drifting lower envelope of the recording while remaining insensitive to transient Ca^2+^ elevations.

The resulting percentile trace was smoothed using a Savitzky-Golay filter to remove high-frequency noise. Any NaN values remaining in the baseline after the rolling operation were corrected by linear interpolation before smoothing. The smoothed baseline was subtracted from the ΔF/F signal to yield a drift-corrected trace.

Following drift correction, a fourth-order Butterworth low-pass filter with a cutoff frequency of 0.3 Hz was applied to the ΔF/F signal to remove high-frequency noise.

### PSTH and trial calcium responses

#### Day 1 – Acquisition

Peri-stimulus time histograms (PSTHs) were constructed for Day 1 acquisition trials by extracting trial-aligned segments of the processed ΔF/F signal. For each light cue onset (t = 0), a window spanning t = -10 s (baseline) to t = +52 s (post-shock) was extracted. This window captured the entire trial structure: pre-light cue baseline (-10 to 0 s), light cue presentation (0 to 20 s), foot shock delivery (20 to 22 s), and a 30 s post-shock period (22 to 52 s).

Response magnitudes were quantified using area under the curve (AUC) calculations applied to predefined time windows: baseline (-10 to 0 s), early light cue (0 to 10 s), late light cue (10 to 20 s), entire light cue (0 to 20 s), and post-shock (20 to 30 s).

For each animal, the trial-averaged ΔF/F signal was first baseline-corrected by subtracting the mean ΔF/F of the pre-cue baseline window (-10 to 0 s) from the entire trial epoch. were then normalized by window duration, facilitating comparisons across windows of different lengths. The trapezoidal rule was used for numerical integration. Trial-by-trial activity was visualized as heatmaps. White horizontal lines delineate individual animal boundaries within the heatmap.

#### Day 3 – Cued test

Ca^2+^ responses during the cued probe test were analyzed by extracting a single trial-aligned segment of the processed ΔF/F signal time-locked to the light cue onset (t = 0). A window spanning t = -10 s (baseline) to t = +60 s (end of cue) was extracted from the full-session Ca^2+^ recording, capturing the pre-cue baseline (-10 to 0 s) and the entire 60 s light cue presentation period (0 to 60 s).

Response magnitudes were quantified using AUC calculations applied to four predefined time windows: baseline (-10 to 0 s), early response (0 to 20 s), late response (20 to 60 s), and full cue response (0 to 60 s). For each animal, AUC values were computed from baseline-corrected ΔF/F traces, where the mean signal of the pre-cue baseline window (-10 to 0 s) was subtracted from the entire epoch prior to integration. AUC values were normalized by window duration to allow comparisons across windows of different lengths. Numerical integration used the trapezoidal rule. Trial-by-trial activity is displayed as heatmaps. White horizontal lines delineated individual animal rows. The light cue onset (t = 0) and, where applicable, cue offset (t = +60 s) were indicated by dashed vertical lines.

### Fiber photometry signal quality control

After analysis of the Day 1 – Acquisition, fiber photometry recordings were subjected to quality control to identify and exclude animals with insufficient signal quality (Simpson *et al*., 2024). First, a signal-to-noise ratio (SNR) was computed as the ratio of the shock-evoked area under the curve to the standard deviation of the ΔF/F signal during a 120 s pre-shock baseline window (-120 to 0 s relative to the first shock). Animals for which this ratio fell below a threshold of 3.0 were flagged for exclusion on SNR grounds. Second, the session dynamic range was computed as the difference between the maximum and minimum ΔF/F values across the entire recording. Animals with a dynamic range below 1.5% were flagged for exclusion on dynamic range grounds. Animals meeting either criterion were excluded from all subsequent analyses.

### Behavioral data processing and freezing detection

#### DeepLabCut tracking and preprocessing

Behavioral tracking data were obtained from DeepLabCut (DLC) video analysis (30 fps), yielding frame-by-frame (x, y) coordinates and likelihood values for three body parts: body center, head, light cue source, and LED. For body center and head tracking points, frames with likelihood below 0.98 were flagged as low-confidence, and their coordinates were replaced via cubic spline interpolation using flanking high-confidence points.

For the light-cue marker, a lower likelihood threshold of 0.95 was applied, as this marker was used exclusively for trial onset detection rather than spatial tracking. Pixel coordinates were converted to centimeters using calibration factors derived from known arena dimensions.

A TTL-triggered red LED was placed within the field of view of the behavioral camera, and its visibility was tracked with a DLC marker. This LED was immediately after the TTL pulse sent to the fiber photometry system, providing a shared temporal reference visible in the video recording for sessions in which no light cue was present (Days 2 Contextual Recall and Context Discrimination).

#### Trial Onset Detection and Temporal Alignment

Day 1 (Acquisition) and Day 3 (Cued Recall), light cue onsets were detected from the frame-by-frame likelihood of the light cue marker exceeding 0.95, with a minimum inter-onset interval of 100 s enforced to prevent spurious re-detection within a single trial. The first detected light cue onset frame defined the behavioral reference time (t = 0). Concurrently, the first TTL 0→1 transition in the Ca^2+^ recording defined the Ca^2+^ reference time (t = 0). All subsequent temporal analyses were expressed relative to these synchronized reference points. The Ca^2+^ fluorescence signal was linearly interpolated to the behavioral frame rate (30 fps) to enable frame-by-frame correspondence between astrocytic and behavioral data.

For Day 2 (Contextual Recall and Context Discrimination), the temporal alignment between the Ca^2+^ recording and the behavioral video was achieved using the TTL-triggered LED. The first frame in which the LED was detected was identified as the LED onset frame, defining the behavioral reference time (t = 0). Concurrently, the first TTL 0→1 transition in the fiber photometry recording defined the Ca^2+^ reference time (t = 0). Both signals were re-zeroed to their respective reference times, aligned to a common timeline, and the initial 10 s following alignment were discarded to remove onset-related transients. The full 3 min session was then analyzed as a single continuous session.

#### Head displacement and angular velocity computation

Inter-frame Euclidean displacement was computed (Pinho *et al*., 2025) from consecutive body center coordinates (in centimeters) as:

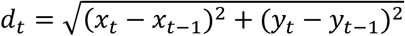

yielding a displacement value in cm per frame. Head orientation at each frame was computed as the arctangent of the vector from the body center to the head position:

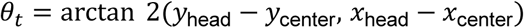

expressed in degrees and mapped to the range [0°, 360°). Frame-to-frame angular differences were computed and corrected for circular wrapping by constraining differences to the range [-180°, +180°]. Angular velocity (° s^-1^) was obtained by multiplying the absolute frame-to-frame angular difference by the frame rate (30 fps). Both displacement and angular velocity signals were smoothed independently using a trailing rolling mean filter with a window of 15 frames (0.5 s at 30 fps).

#### Freezing classification

A frame was classified as a freezing candidate if the smoothed inter-frame displacement fell below 0.02 cm frame^-1^ (locomotion criterion) or the smoothed angular velocity fell below 30°s^-1^ (head movement criterion). These two binary candidate arrays were processed through identical post-processing pipelines before being combined.

Post-processing was applied in the following sequence for each signal independently:

1. Brief state transitions shorter than 15 frames were removed by replacing them with the state value of the immediately preceding frame. This operation is bidirectional: it applies to both short candidate-freezing bouts and short candidate-movement bouts, effectively merging small interruptions into the surrounding dominant state.
2. Following gap-filling, candidate-freezing bouts were required to persist for a minimum of 30 consecutive frames (1 s at 30 fps). Bouts failing this minimum duration criterion were reclassified as non-freezing.

The final binary freezing classification was obtained by requiring that both criteria be satisfied simultaneously: a frame was classified as freezing only if both the locomotion-based and head-movement-based post-processed binary arrays indicated freezing at that frame. This dual-criterion approach reduces false positives arising from transient tracking artifacts while capturing the sustained whole-body immobility characteristic of conditioned fear responses. To validate this automated classification, freezing output was compared against manual scoring performed by an experienced observer using Solomon Coder (version beta 19.08.02; András Péter, Budapest, Hungary) on a subset of videos selected to represent the full range of experimental conditions, including varying ambient light levels and all genotypes included in the study (**Fig. S1a**). High agreement between automated and manual scores confirmed that the dual-criterion thresholds accurately captured behaviorally defined freezing across subjects and recording conditions.

#### Behavioral analysis windows

Freezing metrics (percentage of time spent freezing, number of freezing bouts, total freezing duration) were calculated for the entire CFC session.

For the acquisition session (Day 1), the data were computed within four analysis windows, every 60 s in duration. The baseline window spanned the first 60 s of the 120 s pre-trial recording (t = -120 s to t = -60 s relative to the first light onset). Post-conditioning analysis windows were defined relative to each trial’s light cue onset: the light cue remained on for 20 s, followed by a 2 s shock, giving a shock offset at t = 22 s after light cue onset. Each post-shock analysis window began 5 s after shock offset (t = 27 s after light cue onset) and extended for 60 s (t = 27 s to t = 87 s after light cue onset).

For Day 2 (Contextual Recall and Context Discrimination), freezing metrics were computed across the entire 3 min session (180 s) and analyzed as a single continuous epoch following TTL-based temporal alignment.

For Day 3 (Cued Test), freezing metrics were computed within a single 60 s epoch corresponding to the light cue presentation period (0 to 60 s relative to cue onset), following TTL-luz temporal alignment. This epoch constitutes the sole analysis window for Day 3 behavioral quantification.

#### Freezing onset- and offset-aligned calcium activity

Freezing events previously identified, which respected the criteria of 0.02 cm frame^-1^ (locomotion criterion) and angular velocity below 30° s^-1^ (head movement criterion), were subjected to a validity criterion to avoid overlap.

The freezing events were detected: (1) during the acquisition session (Day 1), for each post-shock analysis window of 60 s; (2) during the contextual recall and discrimination sessions (Day 2), across the entire 3 min sessions; (3) and on the cued test (Day 3) during the 60 s light cue period (0 to 60 s relative to cue onset). Here, a freezing onset (initiation) was accepted only if the preceding non-freezing period and the subsequent freezing period were each ≥ 2 s. A freezing offset (termination) was accepted only if the preceding freezing period and the subsequent non-freezing period were each ≥ 2 s. The duration of each flanking bout was measured from the nearest preceding transition of the opposite type, or from the window boundary when no such transition existed. For each valid event, a peri-event ΔF/F segment spanning -4 to +8 s relative to event time was extracted from the aligned Ca^2+^ time series. Segments containing fewer than five data points within the specified window were discarded.

The rate of change of ΔF/F was quantified by fitting a linear regression to each animal’s mean trace over two peri-event time windows: the pre-event window (-2 to 0 s) and the post-event window (0 to +2 s). Regression was performed on the subset of interpolated time points falling within each window boundary. The resulting slope (ΔF/F % s^-1^) was computed for freezing onset and offset events, for each animal and analysis window.

### Ridge regression decoder

To test whether astrocytic Ca^2+^ responses recorded during fear conditioning predict the strength of contextual fear memory 24 h later, a Ridge regression decoder was implemented and evaluated using leave-one-out cross-validation (LOOCV), with significance assessed via a nonparametric permutation test.

#### Ridge regression model

For each animal, the decoder used the mean AUC per minute in the 20-30 s window following the 3^rd^ foot shock onset. The target variable was the percentage of time spent freezing during Day 2 in Ctx A and Ctx B. Before model fitting, the feature was standardized to a zero mean and unit variance within each LOOCV fold using parameters estimated exclusively from the training observations, to prevent data leakage.

The decoder was a Ridge regression model, which applies an L2 regularization penalty to the coefficient:

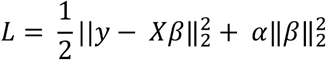

where α controls overall penalty strength. The regularization parameter α was selected automatically via generalized cross-validation (GCV) over a logarithmically spaced grid of 100 values spanning α ∈ [10^-3^, 10^3^] (RidgeCV, scikit-learn).

#### Leave-One-Out Cross-Validation (LOOCV)

Model performance was estimated using LOOCV. For each animal in turn: (i) that animal was removed from the dataset entirely; (ii) a fresh RidgeCV was fitted on the remaining n − 1 animals, including re-fitting the feature scaler and re-selecting α via GCV; (iii) the held-out animal’s standardized feature was transformed using the fold-specific scaler, and its freezing behavior was predicted by the fitted model. After completing all n folds, every animal had exactly one out-of-sample prediction generated by a model that never had access to that animal’s data. Decoding accuracy was quantified as the coefficient of determination R^2^, computed from the n LOOCV predicted-observed pairs. R^2^ = 0 indicates performance equivalent to predicting the group means; R^2^ < 0 indicates performance worse than the mean predictor; R^2^ > 0 indicates good performance.

#### Non-parametric permutation test

The statistical significance of the observed LOOCV R^2^ was assessed by a non-parametric permutation test. The freezing labels were randomly permuted across animals 1,000 times, destroying the real AUC-freezing correlation. For each permutation, the complete LOOCV pipeline was repeated to obtain a null R^2^ value. The permutation p-value was computed as the proportion of permutations yielding an R^2^ equal to or greater than the observed value.

### Open Field Test

The Open Field Test was conducted in white square arenas (30 × 30 × 30 cm) under dim white-light room illumination. Prior to testing, mice were habituated to the recording room for 30 min. Each session lasted 5 min, during which Ca^2+^ fluctuations were recorded via fiber photometry. The arena was cleaned between subjects with 10% ethanol to eliminate olfactory cues.

To visualize the temporal dynamics of Ca^2+^ fluorescence across the full Open Field Test session, ΔF/F traces were displayed as heatmaps for each experimental group. For each animal, the single OFT session trace, extracted and processed as described above, was stored as a single-trial array and loaded accordingly. Traces were arranged as a two-dimensional array with one row per animal and time along the x-axis, spanning the full 300 s session window (*t*_0_ = 0 s, *t*_1_ = 300 s) at 100 Hz. Heatmaps were generated separately for each group. Color scaling was fixed across all animals and groups (minimum: -1 ΔF/F %; maximum: +3 ΔF/F %), with out-of-range values rendered at the respective colormap extremes. Individual animal rows were delineated by white horizontal lines.

Spontaneous Ca^2+^ activity during the Open Field Test was quantified from the continuous ΔF/F trace extracted over the full 5 min session. Spontaneous Ca^2+^ events were detected in the ΔF/F baseline signal using an adaptive threshold approach (Serra *et al*., 2025). A moving standard deviation was computed using a 1 s sliding window. The detection threshold at each time point was set to 2 × the local moving standard deviation, thereby adapting to local signal variability. Peaks in the ΔF/F signal exceeding the adaptive threshold were identified. To exclude noise fluctuations that exceeded the adaptive threshold but lacked biological relevance, only events with an amplitude of at least 1.0 % ΔF/F were kept.

### Histology

#### Euthanasia and brain sectioning

After all procedures, animals were deeply anesthetized with a mixture of ketamine and medetomidine (respectively 150 mg/kg and 0.3 mg/kg, intraperitoneal injection) and transcardially perfused with 0.9% saline, followed by a 4% paraformaldehyde (PFA) solution (pH 7.4 in Phosphate-buffered saline (PBS)). Brains were removed and post-fixed in PFA for 24h (*IP_3_R2 KO* and *WT* mice that were not used for photometry) or removed after whole heads immersion for 48 h in 4% PFA with fiber implant (mice used in photometry). After post-fixation, brains were immersed in 30% sucrose solution at 4°C until sectioned.

Sectioning was performed on a vibrating microtome (VT1000S, Leica Biosystems, Nussloch, Germany) at 40 μm thickness, and the slices were stored in PBS at 4°C.

#### Immunostaining

Sections were washed with PBS solution (3x, 10 mins each), followed by permeabilization with 0.3% Triton-X100 (Sigma Aldrich, USA) in PBS solution (PBS-T) (2x, 10 mins each). The sections were blocked with 10% fetal bovine serum (FBS; in PBS-T at room temperature (RT); Invitrogen, Waltham, MA, USA), to avoid unspecific binding. This blocking step was followed by overnight incubation at 4°C, with the primary AB rabbit polyclonal anti-GFAP (1:200; Dako, Denmark) or AB rabbit polyclonal anti-NeuN (1:200; Cell Signaling, USA), and goat polyclonal anti-GFP (1:400, Abcam, UK) in PBS-T containing 2% FBS.

The following day, sections were washed with PBS-T (3x, 10 mins each) and the secondary AB was added, being incubated at RT for 2 h. Secondaries used were donkey anti-rabbit (1:1000, Alexa Fluor 647, Invitrogen, Waltham, MA, USA) and donkey anti-goat (1:1000, Alexa Fluor 488, Invitrogen, Waltham, MA, USA).

To identify cell nuclei, sections were washed with PBS (2x, 10 mins each), then incubated with 4’,6-diamidino-2-phenylindole (DAPI; 1:1000, 10 mins, RT; Invitrogen, Waltham, MA, USA). At the end, they were mounted on glass slides (SUPERFROST PLUS, Thermo Scientific, Waltham, MA, USA) using Permafluor (Invitrogen, Waltham, MA, USA).

#### Image acquisition and analysis

To assess the localization of virus injection and fiber placement, images from the hippocampus were collected in an inverted fluorescence microscope (Olympus widefield inverted microscope IX81, Tokyo, Japan) using a 20x objective magnification. For cell counting, brain slices where the fiber implant and virus injection were detected were imaged on an Olympus LPS FV1000 confocal microscope (Olympus, Germany) using the 20x (N.A. 0.70; 1.5 μm of z-step; 1024×1024 pixels) objective.

### Statistical analysis

All statistical analyses were performed using GraphPad Prism v8.4.3 (GraphPad Software, San Diego, CA), except where stated otherwise. Data distribution was assessed prior to all comparisons using the Kolmogorov-Smirnov normality test, and the choice of parametric or non-parametric tests was made accordingly.

For comparisons between two groups, normally distributed data were analyzed using a paired or unpaired two-tailed Student’s *t*-test, as appropriate to the experimental design. When normality was not satisfied, the Wilcoxon matched-pairs signed-rank test (paired data) was used instead. Comparisons involving three or more related non-normally distributed groups were performed using the Friedman test, with Dunn’s post hoc correction for multiple comparisons.

For datasets with more than one measurement per subject or condition, a two-way analysis of variance (ANOVA) was applied, followed by Sidak’s post hoc test for multiple comparisons. Where repeated measures were present, but data were incomplete or missing across timepoints, a mixed-effects model was used in place of repeated-measures ANOVA to account for the unbalanced structure of the data, followed by Sidak’s post hoc correction.

Correlations were assessed using Spearman’s rank correlation. All results are presented as mean ± standard error of the mean (SEM). A *p*-value < 0.05 was considered statistically significant.

## Acknowledgments and funding information

The authors thank Leandro A. A. Aguiar (Federal University of Pernambuco, Brazil), Júlia S. Pinho (Gulbenkian Institute for Molecular Medicine, GIMM, Portugal), Marta Navarrete and Javier Sánchez Romero (Cajal Neuroscience Center, CNC, Spain), and the entire C2B team at the ICVS for their helpful suggestions and comments. The authors are grateful to Alfonso Araque (University of Minnesota, USA) and Ju Chen (University of California, San Diego, USA) for sharing the *IP_3_R2 KO* mouse line. The authors acknowledge the Foundation for Science and Technology (FCT) fellowships to DSA and AV; CEECINST/00018/2021 Grant to JFO; grant from “la Caixa” Foundation (LCF/PR/HR21/52410024), Bial Foundation Grant 156/24 to JFO. This work has been funded by national funds, through the Foundation for Science and Technology (FCT), under projects UID/06304/2025 (https://doi.org/10.54499/UID/06304/2025) and LA/P/0050/2020 (https://doi.org/10.54499/LA/P/0050/2020). This work has also been supported by the ICVS Microscopy and Imaging Facility, a member of the Portuguese Platform of Bioimaging (PPBI), funded under project PPBI (POCI-01-0145-FEDER-022122).

## Author contributions

DSA and JFO designed the project. DSA and AV managed the animal colonies, performed the surgeries, and conducted the behavioral experiments. DSA, JFV, and JFO analyzed the results. DSA wrote the Python scripts, which were revised by RMJ. JFV, CSC, AJR, and JFO contributed to the intellectual content and critical manuscript review.

## Ethics declarations

The authors declare no competing interests.

## Supplementary Figure Legends

**Supplementary Figure 1 Freezing quantification and astrocytic responses to the light cue and foot shock.**

**a**, Comparison between the freezing quantified manually and using DeepLabCut (DLC)-based automated analysis (*n* = 17 videos). **b**, AUC analysis of early (0–10 s) and late (10–20 s) Ca^2+^ activity during acquisition (*n* = 14 mice). **c**, AUC analysis of light cue-evoked Ca^2+^ activity across the three conditioning trials (*n* = 14 mice). **d**, AUC analysis of shock-evoked Ca^2+^ across the three conditioning trials (*n* = 14 mice). Data are presented as mean values ± SEM.

**Supplementary Figure 2 Cue-evoked astrocytic responses in the dHIP do not predict contextual fear expression.**

**a**, Correlation between the light cue-evoked Ca^2+^ responses across the three conditioning trials and freezing in Context A (*n* = 14 mice). **b**, Correlation between the light cue-evoked Ca^2+^ responses across the three conditioning trials and freezing in Context B (*n* = 14 mice). Data are presented as mean values ± SEM.

## References

Adamsky A, Kol A, Kreisel T, Doron A, Ozeri-Engelhard N, Melcer T, Refaeli R, Horn H, Regev L, Groysman M, London M & Goshen I (2018). Astrocytic Activation Generates De Novo Neuronal Potentiation and Memory Enhancement. Cell 174, 59–71.e14.

Bukalo O, O’Sullivan R, Tanisumi Y, Mendez A, Weinholtz C, Zimmerman S, Offenberg V, Carpenter O, Bhagwat H, Mosley S, O’Malley JJ, Lyons K, Fang Y, Goldschlager J, Ostroff LE, Penzo MA, Wake H, Halladay LR & Holmes A (2026). Astrocytes enable amygdala neural representations supporting memory. Nature 652, 434–441.

Delepine C, Shih J, Li K, Gaudeaux P & Sur M (2023). Differential Effects of Astrocyte Manipulations on Learned Motor Behavior and Neuronal Ensembles in the Motor Cortex. J Neurosci 43, 2696–2713.

Deseyve C, Domingues AV, Carvalho TTA, Armada G, Correia R, Vieitas-Gaspar N, Wezik M, Pinto L, Sousa N, Coimbra B, Rodrigues AJ & Soares-Cunha C (2024). Nucleus accumbens neurons dynamically respond to appetitive and aversive associative learning. J Neurochem 168, 312–327.

Dewa K et al. (2025). The astrocytic ensemble acts as a multiday trace to stabilize memory. Nature 648, 146–156.

Fedotova A, Brazhe A, Doronin M, Toptunov D, Pryazhnikov E, Khiroug L, Verkhratsky A & Semyanov A (2023). Dissociation Between Neuronal and Astrocytic Calcium Activity in Response to Locomotion in Mice. Function 4, zqad019.

Franklin KBJ & Paxinos G (2007). The mouse brain in stereotaxic coordinates, 3rd ed. Academic Press. Available at: https://ci.nii.ac.jp/ncid/BB0484642X [Accessed May 13, 2026].

Ghenissa O, Guayasamin M, Ngo K, Duquenne M, Peyrard S, Amilhon B & Murphy-Royal C (2026). Basolateral amygdala astrocytes encode anxiety states. Neuron; DOI: 10.1016/j.neuron.2026.02.038.

Guerra-Gomes S, Cunha-Garcia D, Nascimento DSM, Duarte-Silva S, Loureiro-Campos E, Sardinha VM, Viana JF, Sousa N, Maciel P, Pinto L & Oliveira JF (2021). IP3R2 null mice display a normal acquisition of somatic and neurological development milestones. Eur J Neurosci 54, 5673–5686.

Hirrlinger J & Nimmerjahn A (2022). A perspective on astrocyte regulation of neural circuit function and animal behavior. Glia 70, 1554–1580.

Kol A, Adamsky A, Groysman M, Kreisel T, London M & Goshen I (2020). Astrocytes contribute to remote memory formation by modulating hippocampal–cortical communication during learning. Nat Neurosci 23, 1229–1239.

Li X, Zima AV, Sheikh F, Blatter LA & Chen J (2005). Endothelin-1-induced arrhythmogenic Ca2+ signaling is abolished in atrial myocytes of inositol-1,4,5-trisphosphate(IP3)-receptor type 2-deficient mice. Circ Res 96, 1274–1281.

Li Y, Li L, Wang Y, Li X, Ding X, Li L, Fei F, Zheng Y, Cheng L, Duan S, Parpura V, Wang Y & Chen Z (2025). Cholinergic signaling to CA1 astrocytes controls fear extinction. Sci Adv 11, eads7191.

Li Y, Li L, Wu J, Zhu Z, Feng X, Qin L, Zhu Y, Sun L, Liu Y, Qiu Z, Duan S & Yu Y-Q (2020). Activation of astrocytes in hippocampus decreases fear memory through adenosine A1 receptors ed. McCarthy MM, Colgin LL, McCarthy MM & Wu L-J. eLife 9, e57155.

Maren S, Phan KL & Liberzon I (2013). The contextual brain: implications for fear conditioning, extinction and psychopathology. Nat Rev Neurosci 14, 417–428.

Miguel-Quesada C, Zaforas M, Herrera-Pérez S, Lines J, Fernández-López E, Alonso-Calviño E, Ardaya M, Soria FN, Araque A, Aguilar J & Rosa JM (2023). Astrocytes adjust the dynamic range of cortical network activity to control modality-specific sensory information processing. Cell Rep 42, 112950.

Murphy-Royal C, Ching S & Papouin T (2023). A conceptual framework for astrocyte function. Nat Neurosci 26, 1848–1856.

Nagai J, Yu X, Papouin T, Cheong E, Freeman MR, Monk KR, Hastings MH, Haydon PG, Rowitch D, Shaham S & Khakh BS (2021). Behaviorally consequential astrocytic regulation of neural circuits. Neuron 109, 576–596.

Navarrete M, Perea G, Sevilla DF de, Gómez-Gonzalo M, Núñez A, Martín ED & Araque A (2012). Astrocytes Mediate In Vivo Cholinergic-Induced Synaptic Plasticity. PLOS Biol 10, e1001259.

Oliveira JF, Sardinha VM, Guerra-Gomes S, Araque A & Sousa N (2015). Do stars govern our actions? Astrocyte involvement in rodent behavior. Trends Neurosci 38, 535–549.

Petravicz J, Fiacco TA & McCarthy KD (2008). Loss of IP3 Receptor-Dependent Ca2+ Increases in Hippocampal Astrocytes Does Not Affect Baseline CA1 Pyramidal Neuron Synaptic Activity. J Neurosci 28, 4967–4973.

Pinho JS, Ramon-Duaso C, Manzanares-Sierra I & Busquets-Garcia A (2025). Dorsal hippocampus mediates light–tone associations in male mice ed. Holmes N & Wassum KM. eLife 14, RP105863.

Plas SL, Tuna T, Bayer H, Juliano VAL, Sweck SO, Arellano Perez AD, Hassell JE & Maren S (2024). Neural circuits for the adaptive regulation of fear and extinction memory. Front Behav Neurosci; DOI: 10.3389/fnbeh.2024.1352797.

Sánchez Romero J & Navarrete M (2026). Astroengrams: rethinking the cellular substrate for memory. Nat Rev Neurosci 27, 289–300.

Semyanov A, Henneberger C & Agarwal A (2020). Making sense of astrocytic calcium signals — from acquisition to interpretation. Nat Rev Neurosci 21, 551–564.

Serra I, Martín-Monteagudo C, Sánchez Romero J, Quintanilla JP, Ganchala D, Arevalo M-A, García-Marqués J & Navarrete M (2025). Astrocyte ensembles manipulated with AstroLight tune cue-motivated behavior. Nat Neurosci 28, 616–626.

Sherathiya VN, Schaid MD, Seiler JL, Lopez GC & Lerner TN (2021). GuPPy, a Python toolbox for the analysis of fiber photometry data. Sci Rep 11, 24212.

Simpson EH, Akam T, Patriarchi T, Blanco-Pozo M, Burgeno LM, Mohebi A, Cragg SJ & Walton ME (2024). Lights, fiber, action! A primer on *in vivo* fiber photometry. Neuron 112, 718–739.

Srinivasan R, Huang BS, Venugopal S, Johnston AD, Chai H, Zeng H, Golshani P & Khakh BS (2015). Ca(2+) signaling in astrocytes from Ip3r2(-/-) mice in brain slices and during startle responses in vivo. Nat Neurosci 18, 708–717.

Suthard RL, Senne RA, Buzharsky MD, Diep AH, Pyo AY & Ramirez S (2024). Engram reactivation mimics cellular signatures of fear. Cell Rep 43, 113850.

Suthard RL, Senne RA, Buzharsky MD, Pyo AY, Dorst KE, Diep AH, Cole RH & Ramirez S (2023). Basolateral Amygdala Astrocytes Are Engaged by the Acquisition and Expression of a Contextual Fear Memory. J Neurosci 43, 4997–5013.

Tovote P, Fadok JP & Lüthi A (2015). Neuronal circuits for fear and anxiety. Nat Rev Neurosci 16, 317–331.

Veiga A, Abreu DS, Dias JD, Azenha P, Barsanti S & Oliveira JF (2025). Calcium-Dependent Signaling in Astrocytes: Downstream Mechanisms and Implications for Cognition. J Neurochem 169, e70019.

Williamson MR, Kwon W, Woo J, Ko Y, Maleki E, Yu K, Murali S, Sardar D & Deneen B (2025). Learning-associated astrocyte ensembles regulate memory recall. Nature 637, 478–486.

