## Supplementary figures and images for "Hippocampal astrocytes contribute to encoding context-specific aversive stimuli to regulate fear-related behavior"

### Fig. S

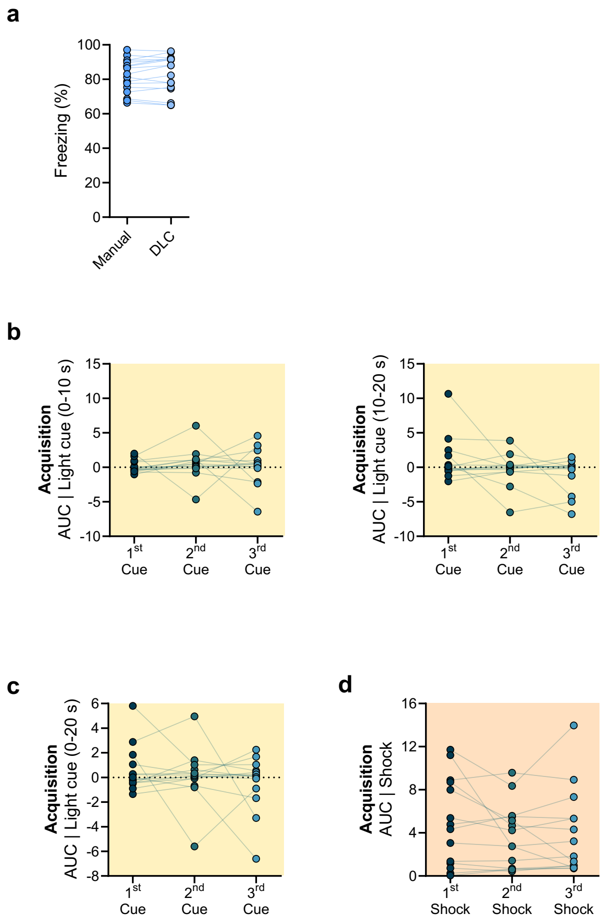

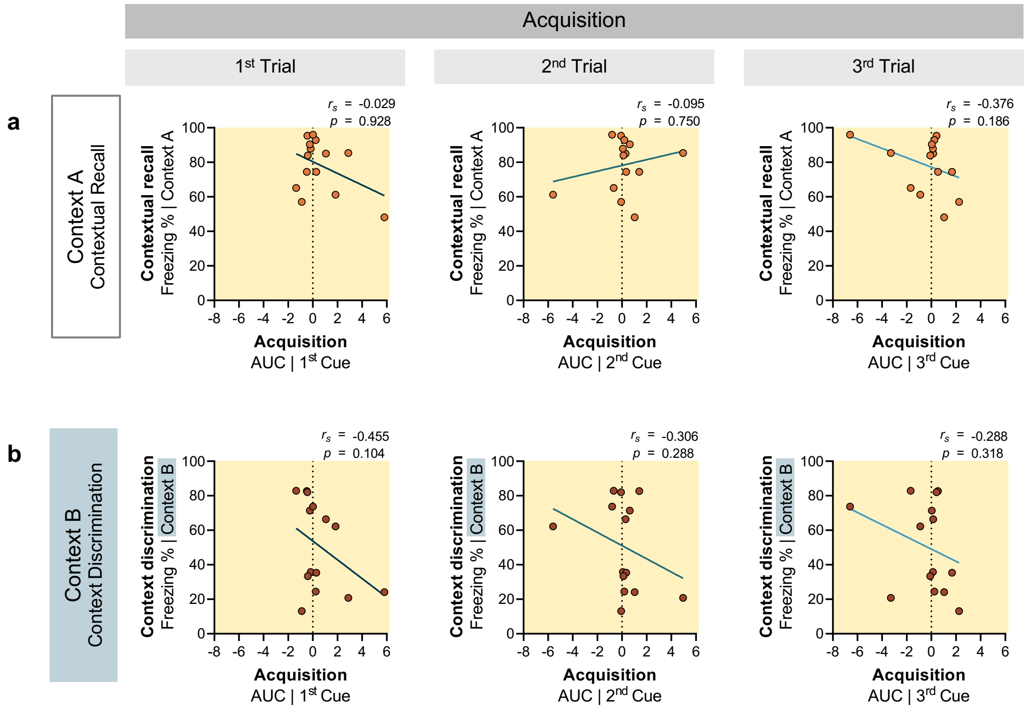
